# Self-Organizing 3D Human Choroid Plexus-Ventricle-Cortical Organoids

**DOI:** 10.1101/2020.09.30.321554

**Authors:** Mohammed R. Shaker, Justin Cooper-White, Ernst J. Wolvetang

## Abstract

Both the choroid plexus (CP) and the cortex are derived from the rostral neural tube during early embryonic development. In addition to producing CSF, the CP secretes essential factors that orchestrate cortical development and later neurogenesis. Previous brain modeling efforts with human pluripotent stem cells (hPSCs) generated either cortical or CP tissues in 3D culture. Here, we used hPSC-derived neuroectodermal cells, the building blocks of the anterior body, to simultaneously generate CP that forms ventricles and cortical cells in organoids (CVCOs), which can be maintained as 3D organoid cultures. Large scale culture revealed reproducibility of the protocol independent of cell lines, clones or batches. CVCOs contain mature and functional CP that projects multiple cilia into the ventricle-like fluid filled cysts and is in direct contact with appropriately patterned cortical cells. CVCOs thus recapitulate key features of developing forebrain structures observed in *in vivo* and constitute a useful for dissecting the role of CP in human forebrain development in health and disease.

## INTRODUCTION

The choroid plexus (CP) is a secretory and a highly vascularized tissue located within each ventricle of the vertebrate brain (Lun et al., 2015). The CP supports the central nervous system by producing cerebrospinal fluid (CSF), producing and transporting a variety of signaling factors that are excreted into the ventricle, and preventing the infiltration of immune cells into the central nervous system (Lun et al., 2015). During early development, shortly after invagination of the forebrain neural ectoderm, the CP anlage and the directly adjacent cortical hem, an important brain organizer region involved in patterning of cortical/hippocampal progenitors and Cajal-Retzius cells, are co-specified and both secrete and respond to morphogens such as Notch, WNT and BMP (Shimogori et al., 2004). This developmental path ensures that the CP is always in close vicinity to the cerebral cortex and this anatomical juxtaposition between CP and the cortex persist throughout life (Emerich et al., 2005). The human CP acquires barrier, secretory and transport capacities after two weeks of development via acquisition of tight junctions and extensive apical microvilli on CP epithelial cells, and influences the fate and migration of neural stem cells and Cajal-Retzius cells of the developing cortex (Lun et al., 2015).

To date mammalian CP development has been predominantly studied in animal models (Zhu et al., 2018, Oshio et al., 2005, Johansson, 2014) and it remains largely unclear to what extent developmental processes are conserved in human or how maldevelopment of the CP contributes to human neurodevelopmental diseases. To model human CP, several 3D culture systems have been developed that allow the generation of CP-like structures *in vitro* starting from human ESC-derived neuroepithelial cells (Watanabe et al., 2012, Sakaguchi et al., 2015, Pellegrini et al., 2020). These models were however not designed to also contain developing neuronal compartments.

Here we report the generation of human cortical brain organoids that are both surrounded by CP <u>and</u> contain properly patterned and developing cortical cell types, providing a novel model for studying the impact of the CP on normal and abnormal early human brain developmental processes.

## RESULTS

### Self-Assembly of Choroid Plexus Organoids Recapitulates Embryonic Development

We used human ES and iPSC lines (Figure S1A) (H9, WTC and G22) to generate human neuroectodermal (hNEct) cells by recapitulating early embryonic developmental events *in vitro* (Caronia-Brown et al., 2014). hNEct are primed to develop exclusively into tissues of the anterior body (Caronia-Brown et al., 2014) and are fated to form the cortex dorsally and the cortical hem ventrally, which then further differentiates into choroid plexus (CP) without generating endodermal and mesodermal derivatives. We therefore reasoned that hNEct cells would be an appropriate starting population for the generation of self-organized cortical organoids surrounded by ventricular structures derived from the CP (here termed CP-Ventricle-Cortical organoids (CVCOs)). Initial exposure of human pluripotent stem cells to dual SMAD inhibitors SB and LDN for 3 days resulted in the efficient generation of hNEct cells characterized by the expression of SOX2, PAX6 and NESTIN (Figure S1B). hNEct cells were next lifted to form 3D aggregates on ultra-low attachment 6-well plates in N2 medium supplemented with bFGF, resulting in spheres formation with a neuroepithelium layer (Figure 1A, day 7) and multiple rosettes at the center expressing SOX2 (Figure S1C). To mimic the secretion of BMP4 and WNTs by the cortical hem that specifies the dorsal cortical hem into CP (Dziegielewska et al., 2001, Watanabe et al., 2012) we treated these spheres with BMP4 and Chiron daily every two days to promote CP formation. Over the next 14 days we observed that organoids gradually increased in size (Figure 1B) with a sharp expansion on day 21, indicative of the differentiation and expansion stages, respectively, until reaching a final a mean core diameter size of 1.2 mm on day 28, that did not further increase over the subsequent week (day 35) (Figure 1B). To reveal the progression of cell fate choices over time we quantified mRNA levels of cortical hem and CP markers by quantitative PCR (qPCR). This showed that 14 to 21 days of Chiron and BMP4 treatment resulted in a significant induction of MSX1/2 expression followed by sharp reduction on day 28 (Figure 1C), suggesting the induction of cortical hem. In agreement with these data cross-sectioned organoids revealed co-expression of cortical hem markers MSX1/2 and LMX1A proteins in the folded epithelial by day 14 (Figure 1D). On day 21 to day 28, we detected the emergence of thin epithelial layers surrounding the organoid (Figure 1A), and qPCR demonstrated the concomitant upregulation of CP markers, AQP1, TTR and KLOTHO (Figure 1C). Consistent with these observations, immunostaining revealed the expression of the definitive CP markers TTR and LMX1A proteins in these epithelial layers (Figure 1D). To better understand the overall tissue architecture of these organoids we performed high-resolution 3D imaging and identified multiple CPs emerging from a single organoid that formed an intact epithelial covering of the entire organoid as indicated by the tight junction marker ZO-1 (Figure 1E). These cells also robustly expressed KLOTHO, an anti-aging protein known to be expressed in mouse CP (Zhu et al., 2018), confirming that KLOTHO expression in the CP is evolutionary conserved and suggesting that its role in these cells can be studied with these human CP organoids. Collectively, these data outline a rapid protocol for generating human brain organoids that are fully encased in CP.

**Figure 1.**
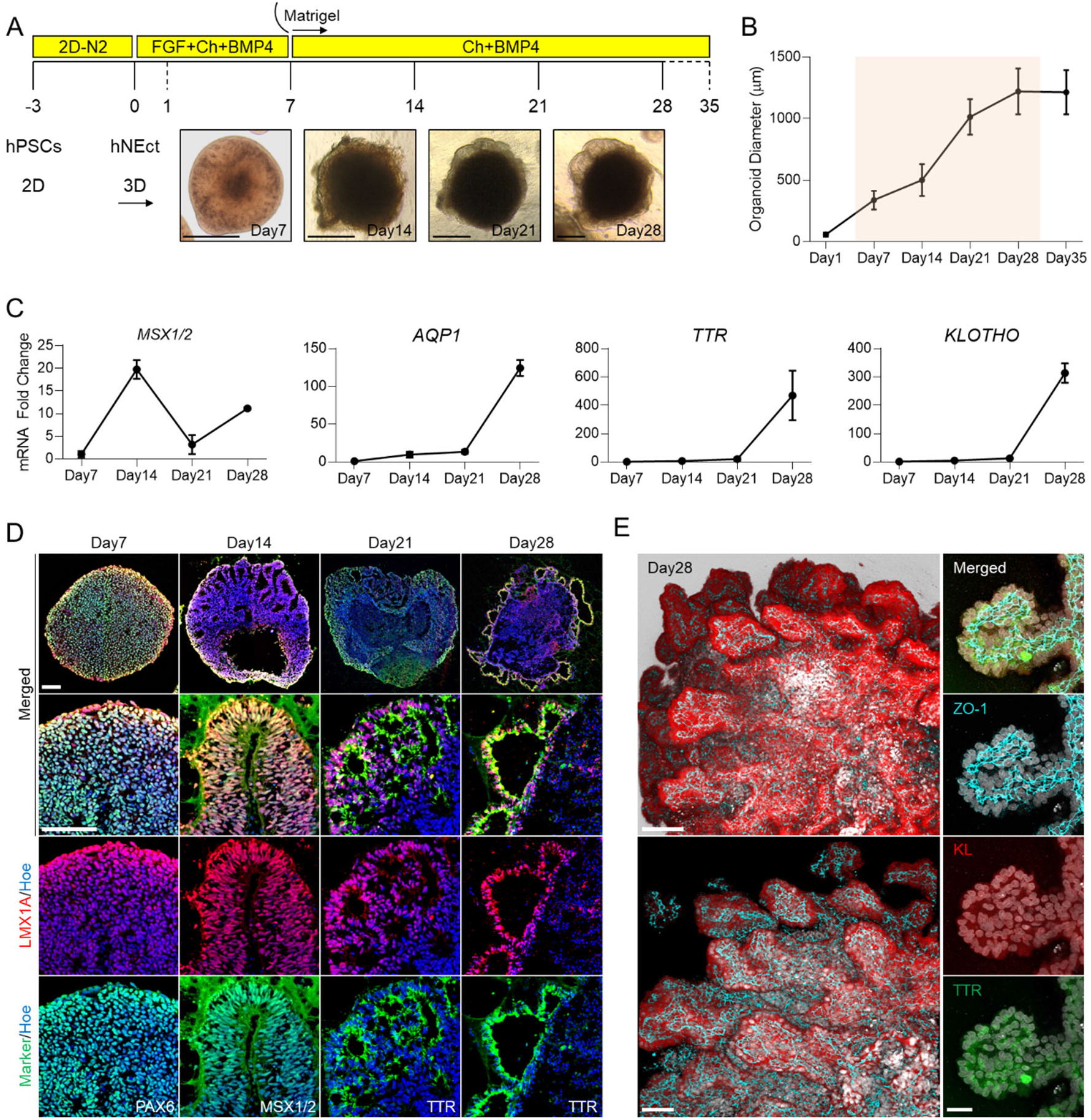
Generation of Human Self-organizing multiple CVCOs in 3D from Neuroectoderm. (**A**) Schematic representation of the strategy used to generate CVCOs from hPSCs. Below brightfield images showing the developmental stages of CVCO overtime *in vitro*. 125 μm. (**B**) Graph showing the growth (average diameter) of CVCOs at different stages of *in vitro* culture. Data are presented as mean ± standard error mean (n = 4). Shadow orange indicates *in vitro* days of CVCOs to be used for subsequent analysis. (**C**) qRT-PCR of cortical hem marker (*MSX1/2*) and CPs definitive markers (*AQP1*, *TTR* and *KLOTHO*). All values were normalized to GAPDH levels of their respective samples and expressed relative to Day 7 values to obtain the fold change. Data are shown as mean ± standard error mean; Number of independent experiments = 4. (**D**) Analysis of immunostaining of sections organoids at days 7, 14, 21 and 28 showing the proteins expression of neuroectoderm markers PAX6 (Green) and LMX1A (Red), cortical hem markers MSX1/2 (Green) and LMX1A (Red), and CP definitive markers TTR (Green), LMX1A (Red). All sections were counterstained with Hoechst 33342 (Blue). Scale bar = 72 μm. (**E**) Wholemount immunostaining images of CVCO at day 28 of differentiation. Left images showing the multiple CPs formation at the periphery of an organoid stained with KLOTHO (Red) and ZO-1 (Cyan), Scale bar = 110 μm. Right images are 100X magnification of a single CP stained with ZO-1 (Cyan), KLOTHO (Red) and TTR (Green), Scale bar = 30 μm.

To exemplify the reproducibility of the system, we quantified the number of organoids that showed the characteristic thin TTR^+^ epithelial layers masking the organoids at day 28 from different hPSC lines (Figures S2A, S2B), and found more than 77% of the organoids exhibited the characteristic merging thin epithelium around the organoids, independent of cell line, clone, or batch (Figure S2C). Collectively our demonstrate that this protocol recapitulates the *in vivo* developmental stages of cortical hem patterning and CP tissue formation.

### Ciliogenesis in CVC-Organoids

Next, we determined whether these self-organized CP layers in CVCOs exhibit similar properties to the *in vivo* CP, and examined the occurrence of events such as the establishment of tight junctions, abundance of mitochondria, polarization and ciliogenesis (Liddelow, 2015). We first examined the formation of tight junctions which are known to form during the early stages of CP development *in vivo* (Lun et al., 2015). qPCR analysis of several tight junction genes revealed a gradual increase of some tight junction genes such as *CLDN11*, *CLDN12* and *PCDH18* (Figure 2A), while others, such as *CLDN2* and *CDH5* were only significantly increased by day 28 (Figure 2A). Consistent with the widespread and contiguous expression of the tight junction marker ZO-1 in the CP layers (Figure 2B), transmission electron microscopy (TEM) confirmed the formation of tight junctions as well as the extensive accumulation of mitochondria typical of CP cells (Figure 2C). The establishment of tight junction is also critical for CP cells to join together to polarize with distinct basal components (Redzic, 2011). We then examined whether the CP cells of CVCOs exhibited apicobasal polarity of membrane proteins critical for normal CP epithelial cell function. Labeling CVCOs with ZO-1 and LAMININ, which accumulate on the apical and basal side of polarized CP epithelial, respectively, revealed the correct formation of membrane basal and apical polarity (Figure 2D, E). Taken together, the establishment of tight junction and apicobasal polarity in CVCOs, as well as the increase of mitochondria in CP that is required for CP homeostasis, cerebrospinal fluid secretion, and neurotrophic factor transport and barrier function (Cornford et al., 1997), suggests that these CPs in CVCOs would have the potential for CSF production and effective barrier formation.

**Figure 2.**
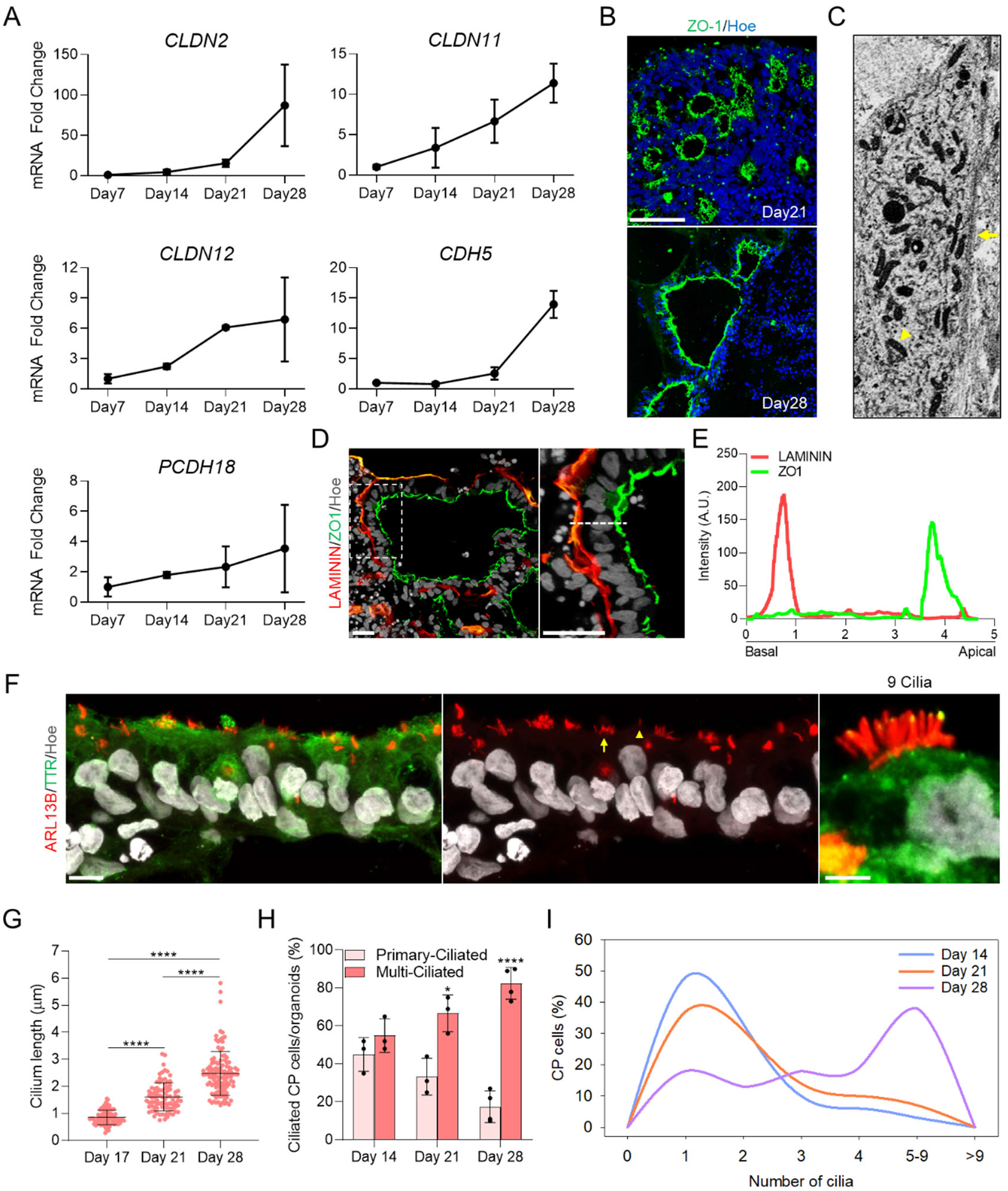
Development and Functional Analysis of CP in CVCOs. (**A**) qRT-PCR of tight junction genes (*CLDN2, CLDN11, CLDN12, PCDH18 and CDH5*) in CVCOs. All values were normalized to GAPDH levels of their respective samples and expressed relative to Day 7 values to obtain the fold change. Data are shown as mean ± standard error mean; Number of independent experiments = 4. (**B**) Analysis of immunostaining of sections CVCOs at days 21 and 28 showing tight junction ZO-1 (Green) protein expression in CPs. All sections were counterstained with Hoechst 33342 (Blue). Scale bar = 72 μm. (**C**) Transmission electron microscopy of CVCOs at day 28 showing the high-density of mitochondria (yellow arrowhead) and tight junction formation (yellow arrow). (**D**) Analysis of immunostaining of sections CVCOs at days 14, 21 and 28 showing the established polarity of basal marked with LAMININ (Red) and apical marked with ZO-1 (Green) proteins expression in CP. All sections were counterstained with Hoechst 33342 (Grey). Scale bar = 20 μm. (**E**) Graph showing the average intensity of LAMININ and ZO-1 expression along the apicobasal of CP epithelium. (**F**) Analysis of immunostaining of sections CVCOs at day 28 showing ARL13B (Red) protein expression in cilia of CP marked with TTR (Green). All sections were counterstained with Hoechst 33342 (Grey). Scale bar = 20 μm., magnified image scale bar = 5 μm. Yellow arrow indicates multi-ciliated cell, yellow arrowhead indicates primary-ciliated cell. (**G**) Quantification of cilia length in CVCOs at day 14, day 21 and day 28 of differentiation Data are presented as mean ± standard deviation; **** P<0.0001 via One Way ANOVA Number of independent experiments = 3. Individual dots represent a cilium length. (**H**) Bar graph demonstrates the percentage of cells with a single cilium and multiple cilia in human CP cells in CVCOs at day 14, day 21 and day 28 of differentiation. Data are presented as mean ± standard deviation; * P<0.05, * P<0.0001 via Student’s t-test. Number of independent experiments = 3. (**I**) The distribution of cilia length in CVCOs culture for day 14, day 21 and day 28. The data is presented as a percentage of CP cells with a single primary cilium or multi-cilia.

Primary cilia in the CP are essential for regulating the flow and transport of CSF and can be characterized according to their length, motility and number per cell (Narita and Takeda, 2015). *In vivo* the CP mainly consists of epithelial cells that are multi-ciliated, with tufts of cilia ranging from 4-8 cilia per cells, but also contains a small fraction of CP cells with one primary cilium extending into the CSF (Lehtinen et al., 2013), suggested to play a role in chemo- and/or osmo-sensation. How and when these cell types are specified in human embryos remains largely unclear. Since the generation of CVCOs closely mimics the progressive morphogenic and temporal sequence of events of normal CP development *in vivo* (Figure 1), we next examined ciliogenesis in human CVCOs cultured for 14, 21 and 28 days. Immunostaining for cilia with the ARL13B antibody demonstrated that human CP cells in CVCOs are ciliated (Figure 2F), and that cilia length significantly increased over time, with a mean value of 0.7μm, 1.6μm and 2.5μm at day 14, day 21 and day 28, respectively (Figure 2G). At day 14 the developing CP contains equal amounts of mono- and multi-ciliated TTR expressing cells, and then displays a progressive increase in multi-ciliated cells to approximately 66% at day 21 and 82% of CP cells at day 28 accompanied by a concomitant decrease in mono-ciliated CP cells (Figure 2H). The number of cilia on CP cells in mammals has been reported to range from 4-8 cilia per cell in rats to 50 cilia per cell in salamander (Lun et al., 2015). Our data show that CP cells in CVCOs display a distinct shift in the number of cilia per cell over time in culture between day 14 and day 28, at which point >30% of all CP cells possess between 5 and 9 cilia per cell (Figure 2I). Collectively these findings indicate that the self-assembled CVCOs recapitulate key aspects of *in vivo* CP ciliogenesis, suggesting that CVCOs may provide a useful model for investigating diseases such as Bardet-Biedl syndrome (Banizs et al., 2005) in which defective primary cilia are thought to cause hydrocephalus.

### Specialized Cellular Compartments of the Cortex Arise in CVCOs

In addition to the mature CPs components outlined above, our organoids are composed of other neural compartments. During embryonic development, the NEct-derived rostral neural tube gives rise to the CP in the dorsomedial telencephalon and cortical plate dorsally (Caronia-Brown et al., 2014). We investigated whether these key cellular compartments of the cortical plate layers arise in our organoids in addition to the CP (Figure 3A) by examining the expression of different cortical neuronal markers known to become specified in cortical organoids (Bhaduri et al., 2020). We found a gradual reduction of *PAX6 mRNA* over time (Figure 3B), and a concomitant increase in layer VI and V neuronal markers as indicated by the expression of *TBR1* and *CTIP2*, respectively (Figure 3B). Staining for these neuronal proteins in midline sectioned organoids revealed specification of cortical TBR1^+^ and CTIP2^+^ neurons in the core of the organoid (Figures 3C, S3). The specification of cortical layer VI and V occurred in close vicinity to the ventricles formed by the CP epithelial cells (Figures 3D, E). This architectural structure of CP, ventricle and cortical cells (Figure 3F) thus closely mimics the developing *in vivo* forebrain structure.

**Figure 3.**
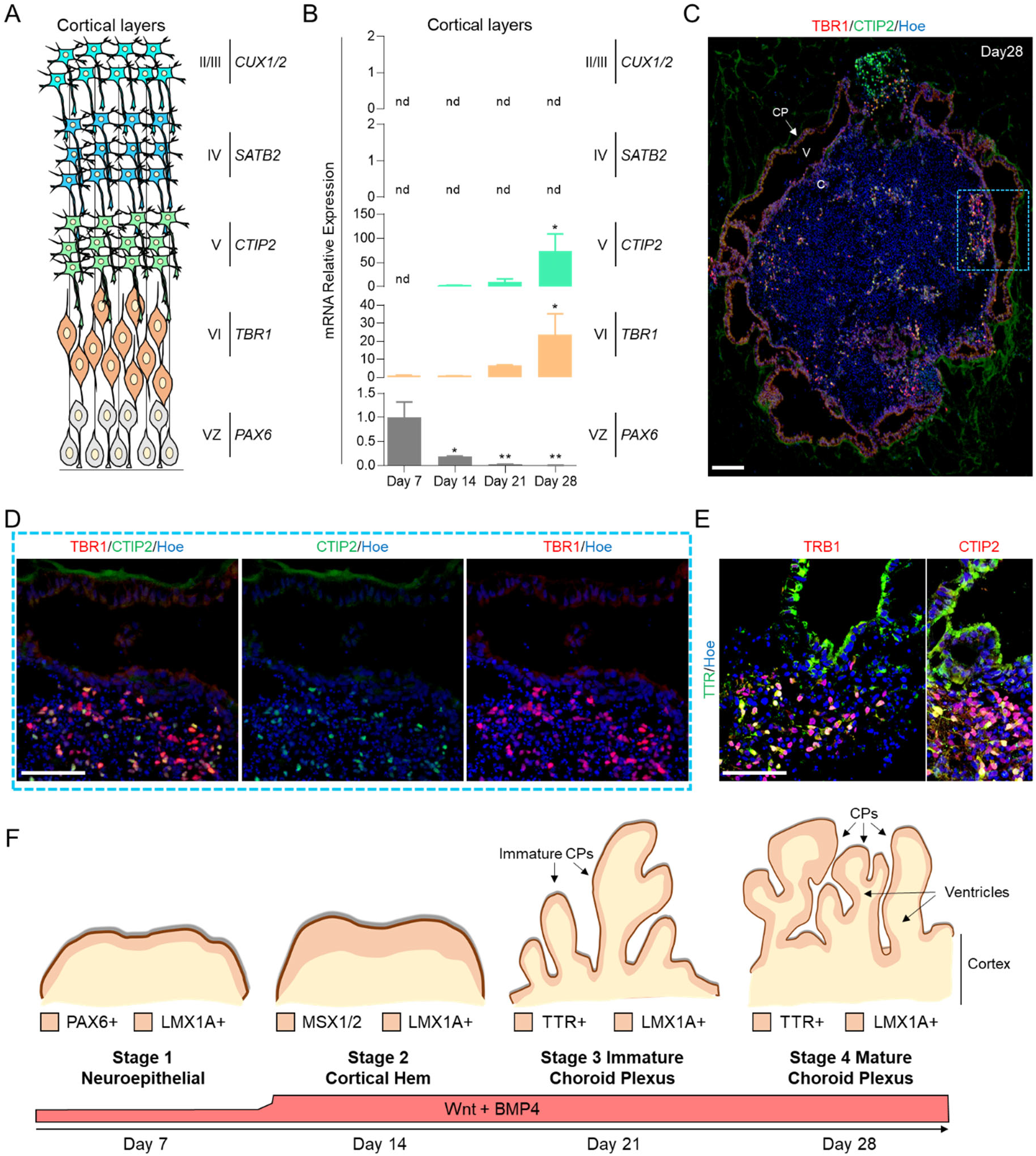
Specialized Cortical neuronal cell types in CVCOs. (**A**) Schematic diagram of the development of the first four cortical plate layers: VZ is ventricle zone. (**B**) qRT-PCR of cortical ventricular zone (VZ) and cortical neuronal layer gene markers (*TBR1*, *CTIP2*, *SATB2* and *CUX1*/2). All values were normalized to GAPDH levels of their respective samples. Data are shown as mean ± standard error mean; Number of independent experiments = 4. nd is not available. (**C**) Analysis of immunostaining of CVCO sections at day 28, showing cortical neurons in layer VI marked by TBR1 (Red) and layer V marked by CTIP2 (Green) proteins. All sections were counterstained with Hoechst 33342 (Blue). Scale bar = 122 μm. (**D**) Magnified images of sectioned CVCOs at day 28 immuno-stained for layer VI cortical neurons marked by TBR1 (Red) and layer V marked by CTIP2 (Green) proteins. All sections were counterstained with Hoechst 33342 (Blue). Scale bar = 72 μm. (**E**) Magnified images of sectioned CVCOs at day 28 immuno-stained for layer VI cortical neurons marked by TBR1 (Red) and choroid plexus marked by TTR (Green), right image is sectioned CVCOs at day 28 immuno-stained with layer V cortical neurons marked by CTIP2 (Red) and choroid plexus marked by TTR (Green). All sections were counterstained with Hoechst 33342 (Blue). Scale bar = 72 μm. (**F**) Schematic diagram to summarize the structure of CVCOs at day 28 showing the specified three domains including CPs, ventricles and the cortex. Below is the schematic condition to induce CVCOs examined in Figures 1, 2 and 3

## DISCUSSION

Congenital disorders encompass a wide range of CP pathologies such as CP cysts, diffuse villous hyperplasia, lipoma and Sturge-Weber syndrome (Naeini et al., 2009). For instance, CP cysts are common in fetuses with trisomy 18 and Aicardi’s syndrome, where CP cysts exhibit accumulation of the CSF causing hydrocephalus (Eide et al., 2020). Many of these CP abnormalities are linked to other structural anomalies in the brain, including cortical development. The development of human *in vitro* models that enable investigation of the cellular and molecular mechanisms underlying such diseases, particularly those in which the early stages of CP development and function are impaired, is therefore valuable, and may allow the development of therapeutic strategies that mitigate disruption of the CP function during development or later in life. In this study, we developed a rapid and robust protocol that generates organoids with multiple CPs that form ventricles that fully enclose developing cortical neuronal cells. We show that the CP epithelial layer that surrounds these organoids exhibits the typical features of the mature *in vivo* CP. Both the cortical cells (progenitors and neurons) and CP cells of these CVCOs arise from a common progenitor, the hNEct cells, that can be readily expanded and genetically manipulated with CRISPR to create various disease models.

While organoids representing multiple brain domains have been established (Grenier et al., 2019, Qian et al., 2019), including the CP (Renner et al., 2017, Pellegrini et al., 2020), the generation of a complex, 3D CVCO from hNEct that contains both CP and cortical cell types has to our knowledge not been reported. Recently, the *in vitro* generation of hNEct from hPSCs has been instrumental in the production of forebrain cells in conventional monolayer culture as well as 3D fashion (Chandrasekaran et al., 2017). Here, we took advantage of the developmental potential of hNEct sheets and generated a complex 3D CVCO model in which two different brain domains, cortical neurons and CP epithelial, developed in parallel from common hNEct progenitors which self-organized, and interacted to promote the formation of ventricle-like structures. Different hPSC lines were used to generate CVCOs, demonstrating the reproducibility of this protocol. The CVCOs contain key components of the mature CP *in vivo* including the establishment of tight junctions and apicobasal polarity. In addition, we also detected high mitochondrial density in CVCO CP epithelium that *in vivo* provides the metabolic capacity for maintaining the secretory activities and ionic gradients across blood-CSF barriers (Cornford et al., 1997), suggesting the CP cells we generated *in vitro* might secrete CSF in a similar fashion to those generated by Pellegrini et al. (Pellegrini et al., 2020).

Previous studies across a number of mammalian species reported the presence of differing amounts of cilia in CP cells (Lun et al., 2015). In this study we show that human CP cells in CVCOs projected up to 9 cilia per cell during differentiation, with a significant shift from primary to multi-cilia during differentiation. Whether these cilia are of the motile 9+2 axoneme kind or non-motile 9+0 axonemal cilia remain to be determined. Previously TEM data demonstrated a 9+2 axoneme in human CPs (Dohrmann and BucY, 1970), while more recent data rather showed a 9+0 axonemal structure in rodents (Narita et al., 2010, Salewski et al., 2012). The CVCOs reported here should provide a useful model to address the axoneme organization and diversity in subsets of human CP cells, and their respective roles in CP function and cortical development (Spassky and Meunier, 2017).

Indeed, the presence of developing cortical cells that are juxtaposed to CP cells in CVCOs has the distinct advantage that it allows investigation of CP mediated neurodevelopmental defects such as hydrocephalus in Bardet-Biedl syndrome through the use of patient specific iPSCs lines or genome edited control iPSC. Because such neurodevelopmental defects can occur very early during embryo development they have been difficult to study in a human setting and our model offers unique opportunities to uncover novel disease processes and to assess the effectiveness of pharmacological interventions designed to rescue CP malformations and promote normal cortical brain development.

Furthermore, it is well established that the CP undergoes a dramatic decline in fluid transport and protein synthesis function during aging and loses its ability to produce and transport growth factors known to regulate neurogenesis (Chen et al., 2012, Narita and Takeda, 2015). With advanced age the epithelial cells of the CP become flattened and cilia are shortened, causing a reduction in the apical surface area (Serot et al., 2003). Many of these CP changes are exaggerated in neurodegenerative diseases such as Alzheimer disease (AD) where an accumulation of β-amyloid in CP epithelial cells was reported to impair CP mitochondrial function and elicit oxidative stress damage (Perez-Gracia et al., 2009). In this study, we showed that the evolutionary conserved anti-aging protein KLOTHO is expressed in human CPs of CVCOs. Klotho is known to protect against multiple neurological and psychological disorders (Vo et al., 2018) and overexpression of Klotho in a mouse model of AD extends lifespan and enhances cognition (Dubal et al., 2015). The CVCOs reported here may thus also enable the elucidation of KLOTHO’s function in the CP and allow testing of its potential to protect CP epithelial and cortical neurons from the accumulation of Aβ in familial AD genetic backgrounds *in vitro* (eg with PS1 mutant iPSC).

The future challenge will be to control anterior-posterior identity of hNEct derived CP while maintaining maturation of cortical neurons. The ability of fusing different domains of the brain using the organoids system has been recently achieved and was used to analyze complex neurodevelopmental defects (Bagley et al., 2017, Xiang et al., 2019). Thus, the establishment of CVCOs with a lateral ventricle identity affords the potential to establish the fourth ventricle CPs with hippocampus, for example through judicious sonic hedgehog activation, and may offer an opportunity to establish and study the anterior-posterior identity with complex networks in an all-brain *in vitro* 3D model in the future.

## MATERIAL AND METHODS

### Human embryonic stem cells culture and cortical organoids generation

Human embryonic stem cells H9 from (Wisconsin International Stem Cell Bank, WiCell Research Institute, WA09 cells), WTC iPSC (gift from Professor Bruce Conklin) and G22 iPSCs lines (available in our lab) gene were cultured according to Stem Cell Technologies protocols that can be found in (https://www.stemcell.com/maintenance-of-human-pluripotent-stem-cells-in-mtesr1.html) on feeder free in hESCs medium on Matrigel (StemCell Technologies, Cat. #354277) in mTeSR (Stem Cell Technologies, Cat. #85851). To generate CVCOs, embryoid bodies were expanded for four days in N2 medium: DMEM/F12 (Gibco, Cat. #11320-33), 2% B-27 supplement (Gibco, Cat. # 17504044), 1% N-2 supplement (Gibco, Cat. #17502-048), 1% MEM Non-Essential Amino Acids (Gibco, Cat. #11140-050), 1% penicillin/streptomycin (Gibco, Cat. #15140148), 0.1% β-mercaptoethanol (Gibco, Cat. #21985-023), embryoid bodies were supplemented daily with (bFGF, 20 ng/mL; R&D, Cat. #233-FB-01M), 3μM Chiron (Sigma Aldrich, Cat. # SML1046-5MG) and 0.5ng/ml of BMP4 (Thermofisher, Cat. # PHC9391). Patterned embryoid bodies were then embedded in Matrigel (StemCell Technologies, Cat. #354277) and switched to the terminal differentiation medium DMEM-F12 (Gibco, Cat. #11320-33): Neurobasal medium (Gibco, Cat. #A35829-01) 1% N2 (Gibco, Cat. #17502-048), 12.5μl of insulin (Sigma), 2% GlutaMAX, 1% MEM Non-Essential Amino Acids (Gibco, Cat. #11140-050), 1% penicillin/streptomycin (Gibco, Cat. #15140148), 0.1% β-mercaptoethanol (Gibco, Cat. #21985-023), and 2% B-27 supplement (Gibco, Cat. # 17504044), supplemented with 3μM Chiron and 50ng/ml of BMP4. Fresh media was replaced three times a week. All experiments were carried out in accordance with the ethical guidelines of the University of Queensland and with the approval by the University of Queensland Human Research Ethics Committee (Approval number-2019000159).

### Immunohistochemistry

Tissue processing was performed as described in (Lee et al., 2019) and immunohistochemistry (IHC) was performed as described in (Shaker et al., 2015). In brief, organoids were fixed in 4% PFA for 60 min at RT, followed by washing with 1x phosphate buffer saline (PBS) three times for 10 min at RT. Fixed organoids were then immersed in 30% sucrose in PBS at 4 °C and allowed to sink before being embedded in a solution containing at 3:2 ratio of Optimal Cutting Temperature (O.C.T) and 30% sucrose on dry ice. Mounted tissues were then subjected to serial sections at 14-μM thickness and collected onto Superfrost slides (Thermo Scientific, cat. #SF41296). To performed IHC, sectioned organoids were washed three times with 1x PBS for 10 minutes at RT before blocking for 1 hour with 3% bovine serum albumin (BSA) (Sigma, Cat. A9418-50G) and 0.1% triton X-100 in 1x PBS. Primary antibodies were added overnight at 4 °C before washing three times with PBS for 10 minutes each at RT. For immunocytochemistry (ICC), cells were allowed to be fixed with 4% PFA in 1x PBS for 10 min at RT. The cells were then washed three times with 1x PBS at RT before blocking and adding primary antibody as stated above. Tissues and cells were then incubated with appropriate secondary antibodies an hour at RT before mounting and imaging. For wholemount, was performed as described in (Yokomizo et al., 2012). All samples were counterstained with Hoechst 33342 (Invitrogen, Cat. #H3570). All images were acquired using confocal microscopy (Leica TCS SP8) based in SBMS Imaging Facilities based at the University of Queensland. The primary antibodies used in this study are listed in Table S1. Alexa-488, Alexa-546, and Alexa-633-conjugated secondary antibodies were obtained from Jackson ImmunoResearch Laboratory.

### qRT-PCR

Total RNA was isolated from organoids as described previously (Lee et al., 2020). For qPCR, 1μg of isolated RNA was utilized to generate the complementary DNA (cDNA) using the first strand cDNA Synthesis Kit (Thermo scientific, Cat. #K1612). SYBR Green (Applied Biosystem, Cat. #A25742) was utilized, and PCR standard reaction conditions were set according to the manufacturer’s instructions. PCR primers were designed using the NCBI free online system and list of primers are listed in Table S2. All experiments were performed in biological triplicates for every sample, and the expression values were normalized against the GAPDH expression value of each sample. The means and standard deviations were calculated and plotted using the GraphPad Prism 8.3.1^®^.

### Transmission Electron Microscopy

Transmission electron microscopy were proceed according to (Hua et al., 2015). In brief, organoids were in 0.1 M Sodium cacodylate buffer in ddH20 containing 2.5% Glutaraldehyde and 2% Paraformaldehyde over night at 4°C. Organoids were first washed three times in 0.1 M Sodium cacodylate buffer for 10 min at RT, followed by immersing in 2% osmium tetroxide (2 ml 4% osmium tetroxide and 2 ml 0.2M cacodylate buffer) for 90 min at RT. The staining buffer was then replaced with 2.5% potassium ferricyanide (4.8 ml 0.2M cacodylate buffer and 4 ml 6% potassium ferricyanide) for 90 min at RT. The organoids were then washed in water three to five times at RT until the water is completely clear, before dipping the organoids in the thiocarbohydrazide solution (0.1g thiocarbohydrazide (locked cupboard) in 10 mL water) for 45 min at 40°C. To remove the background stain, organoids were washed four to five times in water at RT before immersing the organoids into 2% osmium tetroxide in 0.1M cacodylate buffer for 30 min at RT. Organoids were again washed in water to remove the black background. Washed organoids were then dipped into 1% uranyl acetate overnight at 4°C, followed by incubation in 0.03M aspartic acid solution for 30 min at 60°C. Incubated organoids were then washed three times for 30 min at RT. For embedding, the organoids were dehydrated in a series of ethanol 20%, 50%, 60%, 70%, 80%, 90% and 100% for 30 time at RT each. Organoids were then infiltrate with Durcupan resin for 12h at RT. Embedded organoids were then incubated for 48h in 60°C before trimming and imaging.

### Statistical analysis

Normally distributed data were expressed as the mean ± standard deviation of the mean of independent experiments. For non-normally distributed data, the median ± standard deviation was used to express the values. The number of biological replicates as well as the sample size are indicated in the figure legends. The Student’s t-test and one-way or two-way ANOVA were utilized for comparing two and more than two groups, respectively. The Tukey’s post-hoc analysis was applied for comparisons to a single control. Statistical analysis was performed using GraphPad Prism 8.3.1^®^ software. Minimal statistical significance was defined at P < 0.05.

## ACKNOWLEDGMENTS

M.R.S. is supported by the MRFF Leukodystrophy Flagship – Massimo’s Mission (EPCD000034) and by the UQ Centre for Stem Cell Engineering and Regenerative Engineering (UQ stemCARE). This work was funded by UQ Centre for Stem Cell Engineering and Regenerative Engineering (UQ stemCARE). E.W. is supported by the Australian National Health and Medical Research Council (applications 1138795, 1127976, 1144806 and 1130168), BrAshA-T foundation, Perry Cross Spinal Research Foundation and the Australian Research Council. Bruce Conklin (Department of Medicine Gladstone Institute of Cardiovascular Disease) is greatly acknowledged for the WTC iPSCs gift. The authors acknowledge the facilities, and the scientific and technical assistance, of the Microscopy Australia Facility at the Centre for Microscopy and Microanalysis (CMM) of The University of Queensland.

## CONFLICT IF INTEREST

The authors declare no competing interests.

## AUTHOR CONTRIBUTIONS

M.S. performed, analyzed and designed experiments, interpreted the results, and wrote the manuscript. J.C-W contributed to the conception and supervision of the study, interpreted results, and contributed to the writing and editing of the manuscript. E.W. conceived and supervised the study, interpreted results, and wrote the manuscript. The final version of the manuscript was approved by all authors.

## DATA AVAILABILITY STATEMENT

The data that support the findings of this study are available from the corresponding author upon request.

## Supplemental Information

**Figure S1.**
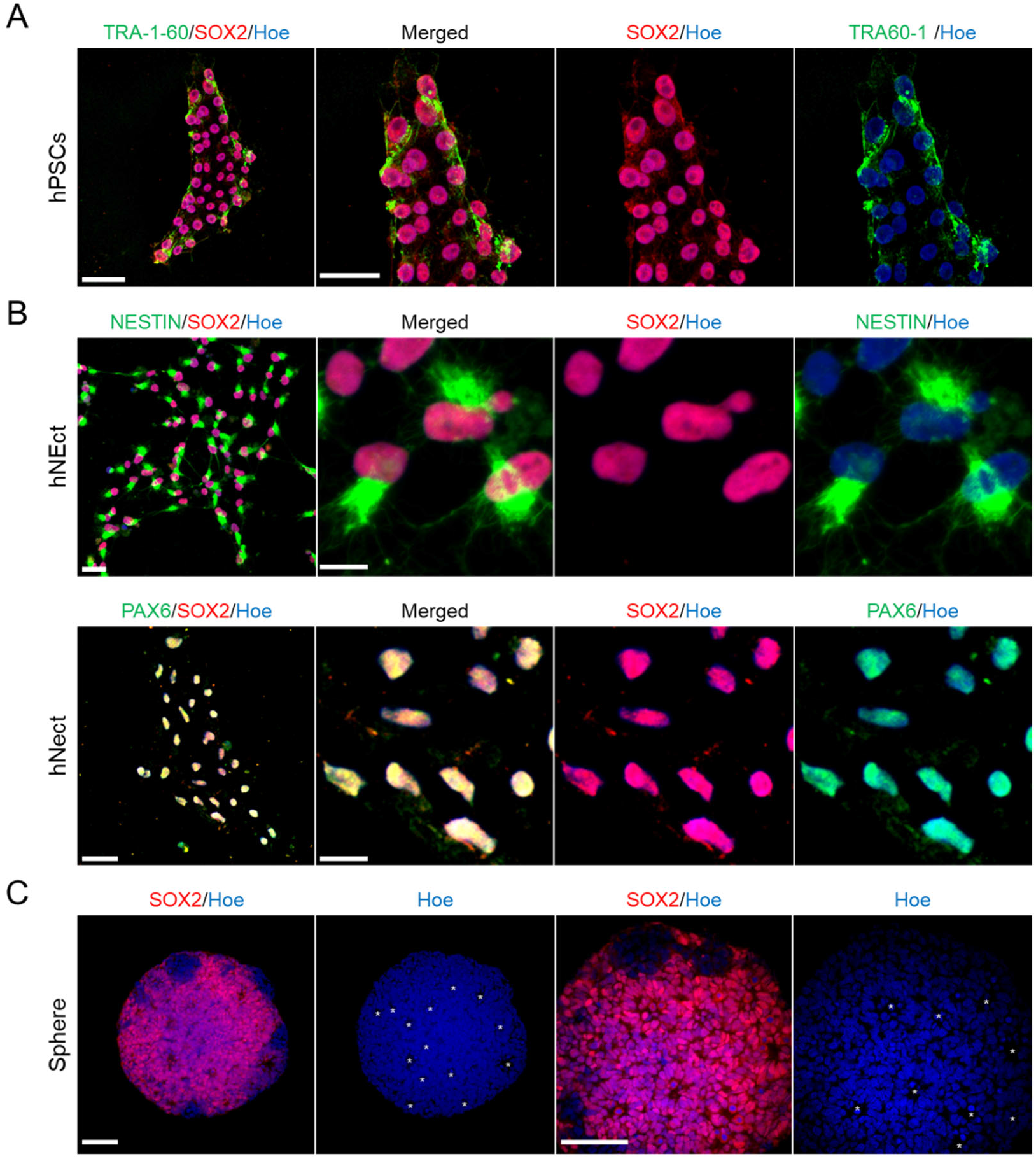
Characterization of Neuroectoderm Differentiation and Spheres Formation from hPSCs, related to Figure 1. (**A**) Immunocytochemistry of human iPSCs stained with TRA-1-60 (Green) and SOX2 (Red) All cells were counterstained with Hoechst 33342 (Blue). Scale bar = 40 μm. (**B**) Immunocytochemistry of human iPSCs-derived neuroectoderm labeled with SOX2 (Red) and NESTIN (Green) antibodies. All cells were counterstained with Hoechst 33342 (Blue/Gray). Scale bar = 20 μm. (**C**) Cross-cut image of a wholemount 3D imaged spheres stained with SOX2 (Red). All cells were counterstained with Hoechst 33342 (Blue/Gray). Scale bar = 60 μm. White stars indicate the multiple rosettes formation.

**Figure S2.**
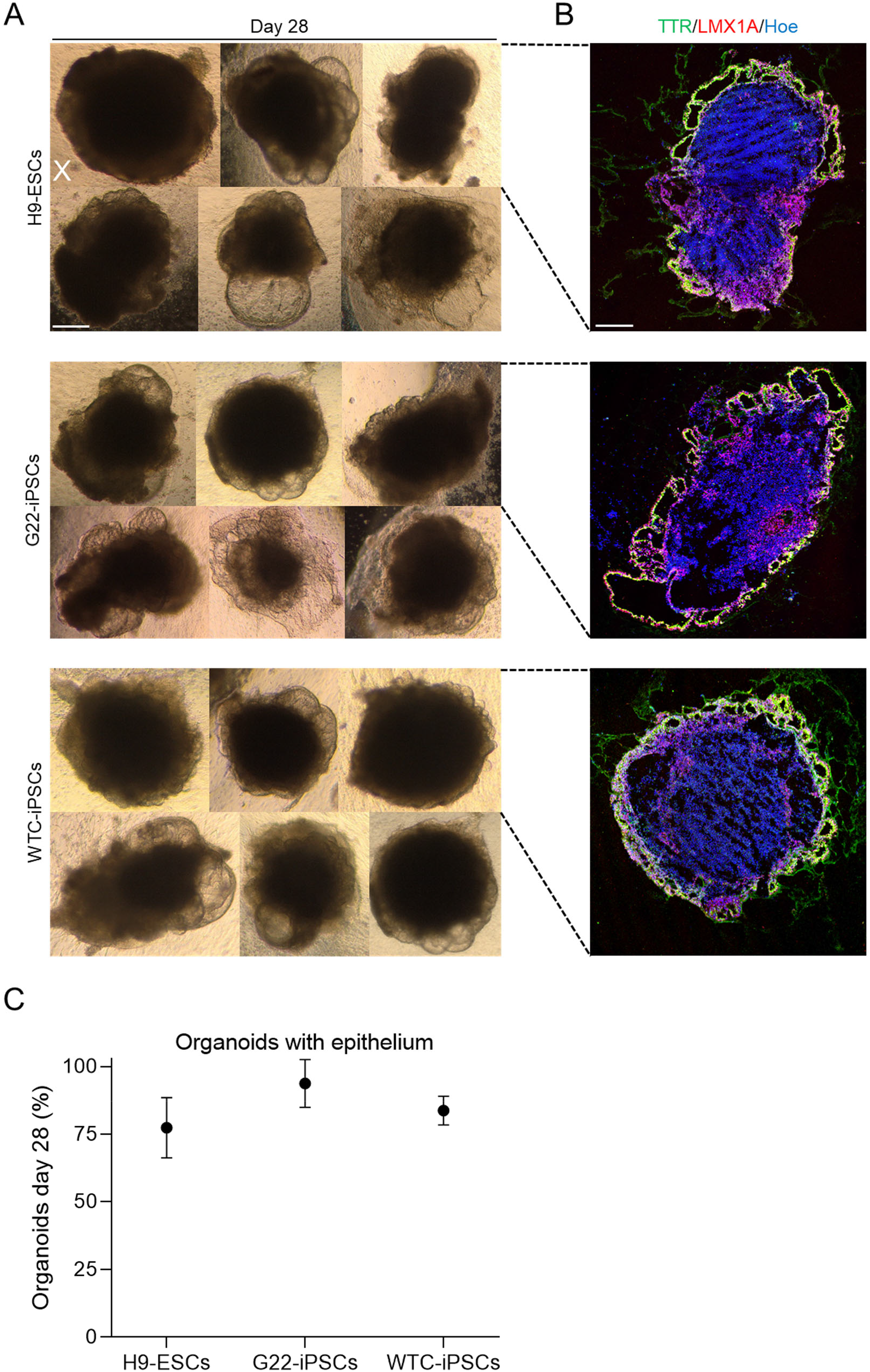
Reproducible Organization of CPs and Cortical Cells in CVCOs Generated from Different hPSC lines, related to Figure 1. (**A**) Bright-field images of CVCOs generated from H9-ESCs, G22-iPSCs and WTC-iPSCs line at day 28 of differentiation. Cross indicate the organoids without emerging thin epithelium. Scale bar = 125 μm. (**B**) Representative images of sectioned CVCOs immune-stained with TTR (Green) and LMX1A (red). All cells were counterstained with Hoechst 33342 (Blue). Scale bar = 200 μm. Dotted black lines indicates the cut bright-field CVCOs. (**C**) Percentages of the successful generation of CVCOs at day 28 in different hPSC lines (H9-ESCs, G22 iPSCs and WTC iPSCs). N = 3. Data are presented as mean ± standard deviation.

**Figure S2.**
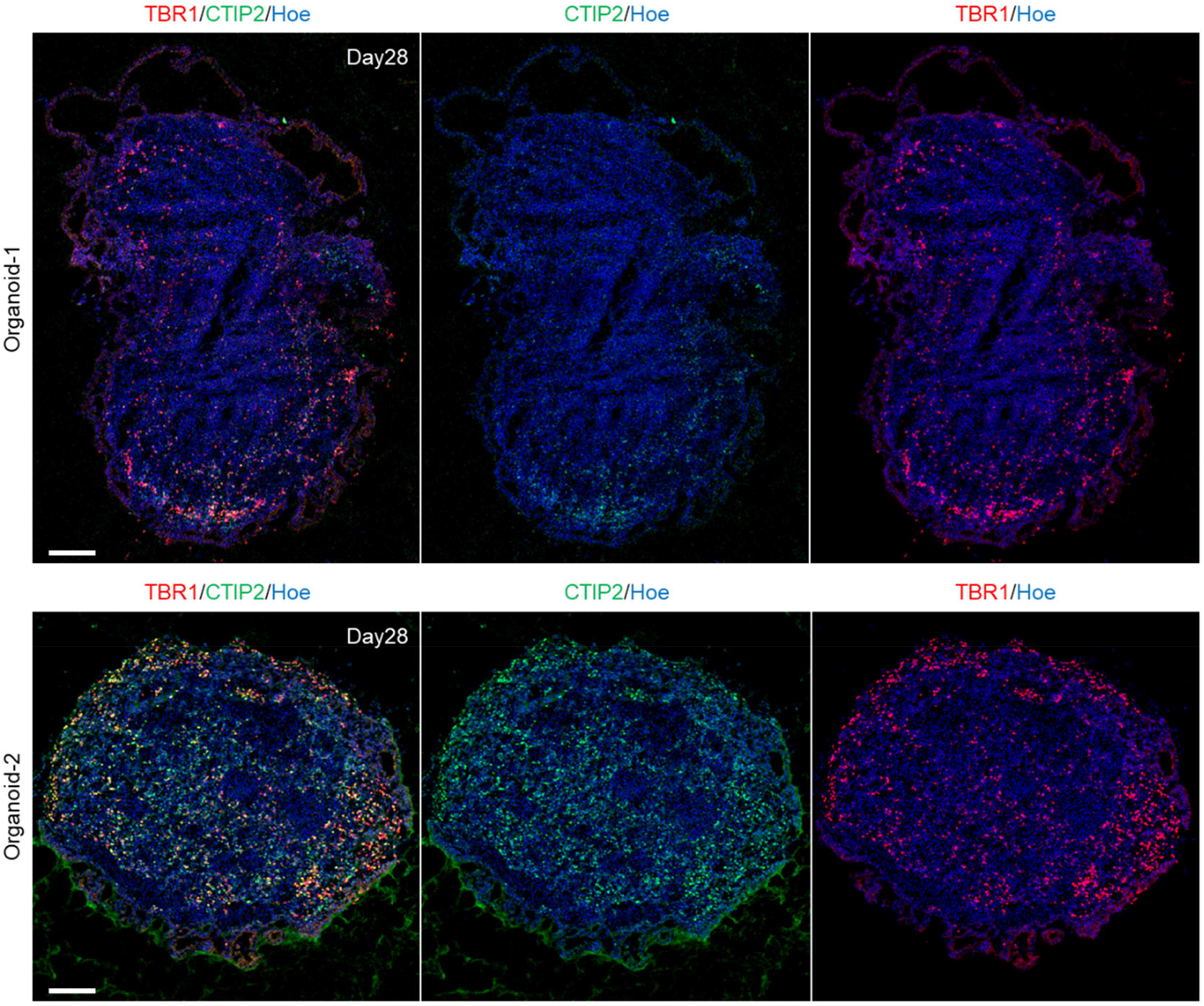
Validation of Cortical Neurons Emerging in CVCOs Generated from Different Batches, related to Figure 3. Analysis of immunostaining of sections CVCOs at day 28 obtained from different batches showing cortical neurons in layer VI marked by TBR1 (Red) and layer V marked by CTIP2 (Green) proteins. All sections were counterstained with Hoechst 33342 (Blue). Scale bar = 122 μm.

**Table S1.**
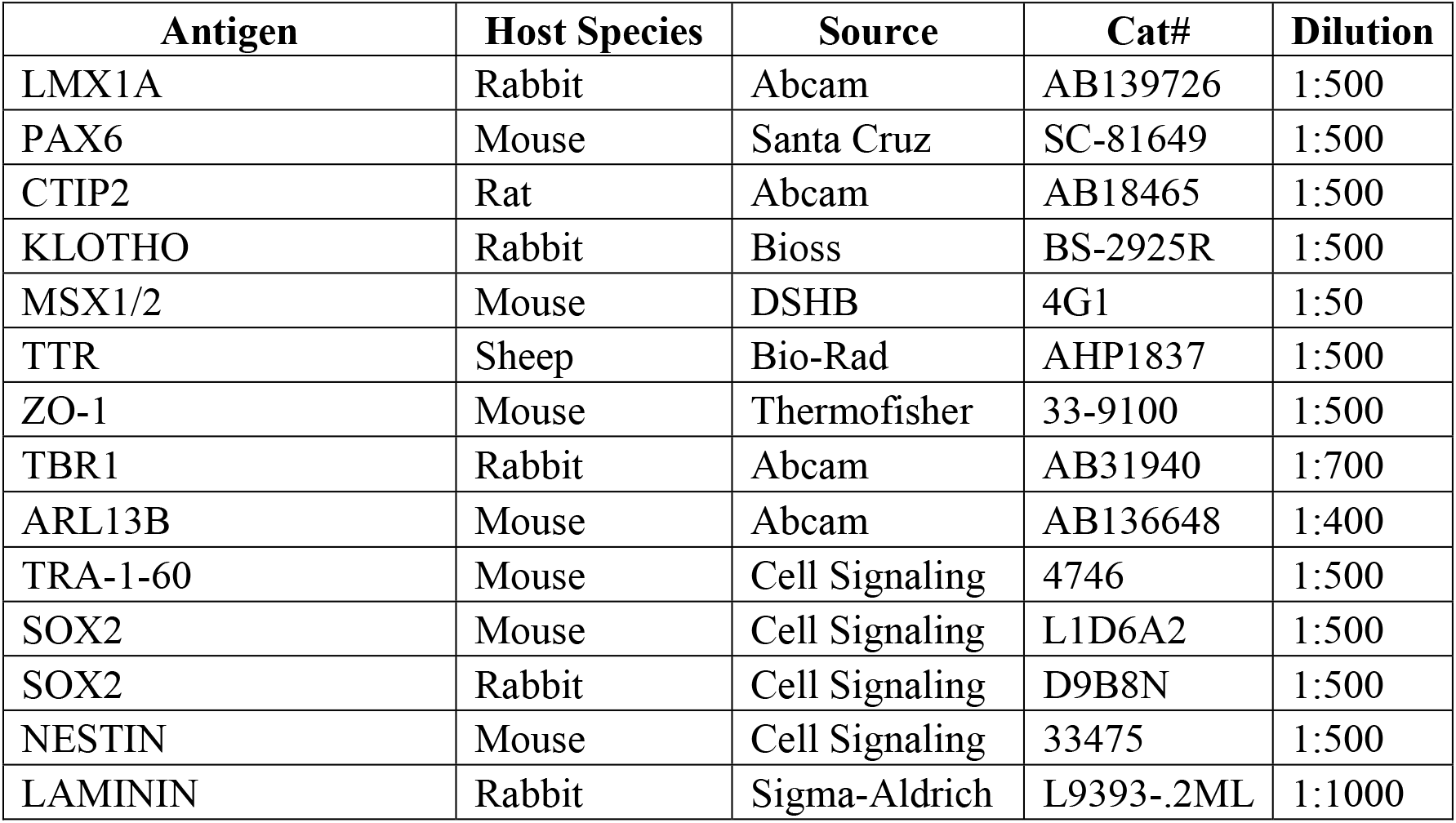
List of Antibodies used for immunohistochemistry

| Antigen | Host Species | Source | Cat# | Dilution |
| --- | --- | --- | --- | --- |
| LMX1A | Rabbit | Abcam | AB139726 | 1:500 |
| PAX6 | Mouse | Santa Cruz | SC-81649 | 1:500 |
| CTIP2 | Rat | Abcam | AB18465 | 1:500 |
| KLOTHO | Rabbit | Bioss | BS-2925R | 1:500 |
| MSX1/2 | Mouse | DSHB | 4G1 | 1:50 |
| TTR | Sheep | Bio-Rad | AHP1837 | 1:500 |
| ZO-1 | Mouse | Thermofisher | 33-9100 | 1:500 |
| TBR1 | Rabbit | Abcam | AB31940 | 1:700 |
| ARL13B | Mouse | Abcam | AB136648 | 1:400 |
| TRA-1-60 | Mouse | Cell Signaling | 4746 | 1:500 |
| SOX2 | Mouse | Cell Signaling | L1D6A2 | 1:500 |
| SOX2 | Rabbit | Cell Signaling | D9B8N | 1:500 |
| NESTIN | Mouse | Cell Signaling | 33475 | 1:500 |
| LAMININ | Rabbit | Sigma-Aldrich | L9393-.2ML | 1:1000 |

**Table S2.**
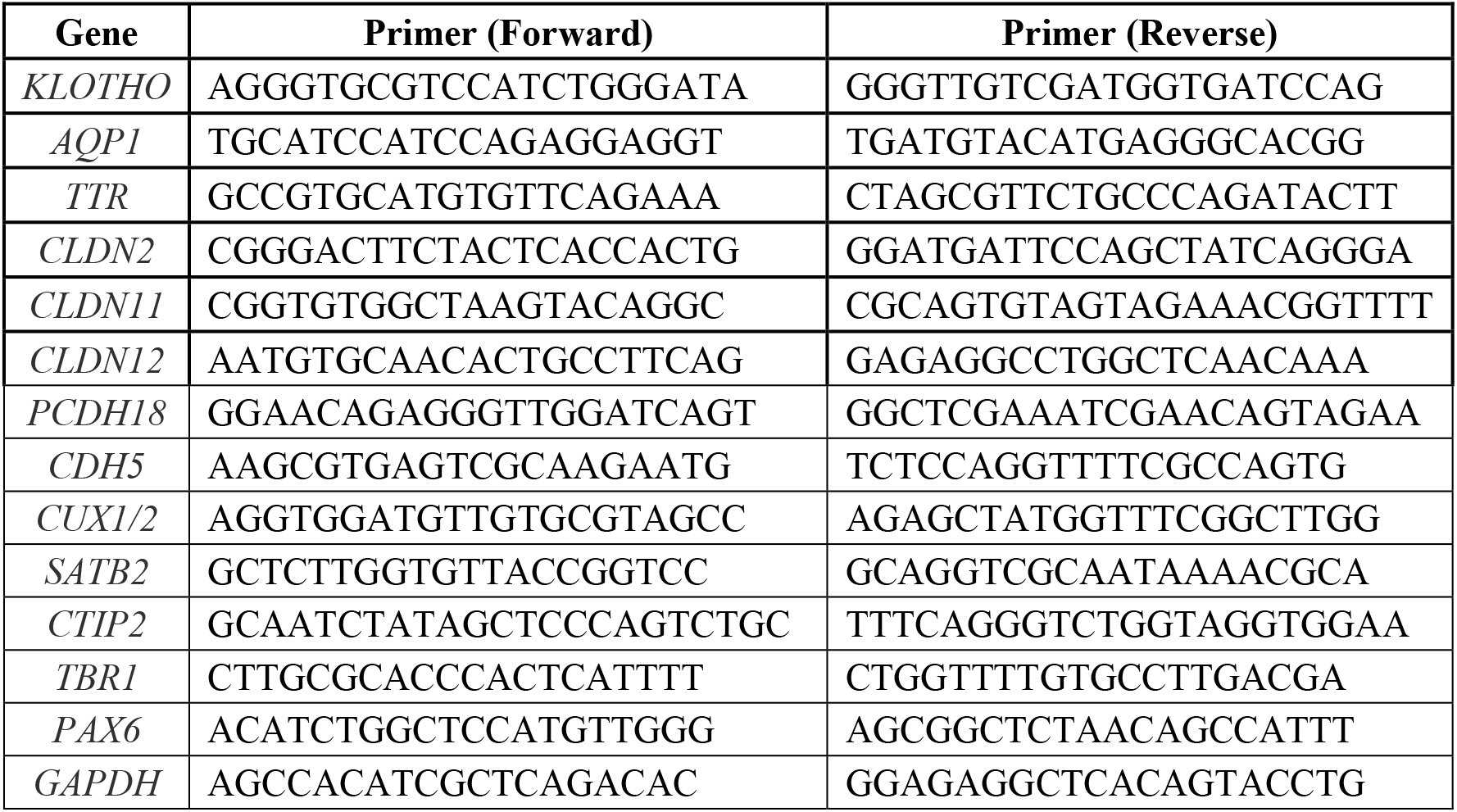
List of Primer Sequences used for RT-PCR (5′−3′ orientation).

